# Coordinated assembly and release of adhesions builds apical junctional belts during *de novo* polarisation of an epithelial tube

**DOI:** 10.1101/2020.02.16.950543

**Authors:** Andrew C Symonds, Clare E Buckley, Charlotte A Williams, Jonathan DW Clarke

## Abstract

Using the zebrafish neural tube as a model, we uncover the *in vivo* mechanisms allowing the generation of two opposing apical epithelial surfaces within the centre of an initially unpolarised, solid organ. We show that NOK/Pals1/Mpp5a and Rab11a play a dual role in coordinating the generation of ipsilateral junctional belts whilst simultaneously releasing contralateral adhesions across the centre of the tissue. We show that Nok and Rab11a mediated resolution of cell-cell adhesions are both necessary for midline lumen opening and contribute to later maintenance of epithelial organisation. We propose these roles for both Nok and Rab11a operate through the transmembrane protein Crumbs. In light of a recent conflicting publication, we also clarify that the junction remodelling role of Nok is not specific to dividing cells.

## Introduction

Epithelia are one of the fundamental tissue types of the body, forming protective sheets of cells around the outside of the organism and lining the inside surface of many organs or parts of organs. All epithelia are polarised and have an apical and a basal surface that are molecularly and functionally distinct. In some cases, the organisation of the apical surface of an epithelium is generated at the free surface of an embryonic sheet of cells while in others the apical surface emerges from within a dense rod or ball of cells to generate an epithelial tube or cyst (Martin-Belmonte et al., 2008, Datta et al., 2011). Generating an apical surface from within a rod or ball of cells adds an extra layer of complexity to this process as cells at the centre of these structures have to lose inappropriate connections to generate a free surface as well as organising that free surface into a typical apical structure comprising a lattice of closely adherent cells connected by apicolateral belts of specialised junctions (typically adherens junctions and tight junctions in vertebrates). A key step, elucidated in epithelial development in flies, seems to be the remodelling of junctional protein location such as Par3 and Cadherin from the developing apical surface to the apico-lateral border in order to build apico-lateral junctional belts (Grawe et al., 1996, Harris & Peifer, 2005, Morais-de-Sa et al., 2010, Tepass, 1996). To what extent similar mechanisms of apical surface development are employed at a free surface compared to within a rod or sphere primordium, and how these can be coordinated with the remodelling of cell-cell connections necessary for *de novo* surface generation, is poorly understood, especially in vertebrates *in vivo*.

Here we analyse the spatial deployment of the cardinal polarity protein Partitioning-defective 3 (Pard3) and the cell adhesion protein N-cadherin (Cdh2) during generation of the apical surface of the neural tube in the zebrafish embryo *in vivo*. In this system the apical surface is generated *de novo* from within the solid neural rod and the relative accessibility and transparency of the zebrafish embryo provides an advantageous system in which to address mechanism in whole embryos. We use experimental manipulations of the NOK/Pals1/Mpp5a scaffold protein and the endocytic recycling protein Rab11a to determine the cellular and molecular mechanisms that simultaneously remodel cell-cell connections in order to release cell adhesions across the organ midline and generate the canonical apical junctional belt organisation of epithelia within a solid primordium. We compare these to previously shown mechanisms that generate apical organisation at a free surface. Our results show that both Mpp5a and Rab11a are required to remodel connections between contralateral and ipsilateral cells and we suggest they operate through apical recruitment of the transmembrane protein Crumbs.

## Results

### Apical rings of Pard3 and ZO-1 are built up from the ventral floor plate

The apical surface of epithelia is characterised by a lattice-like arrangement of polarity and scaffolding proteins (such as Pard3, atypical protein kinase C (aPKC), Zonula occludens-1 (ZO-1)) and cell adhesion proteins (such as Cdh2). We visualised this organisation prior to lumen opening in the sagittal plane of the developing zebrafish neural rod (stages 10 to 18 somites, 14 to 18 hours post fertilization, hpf) at the level of the nascent anterior spinal cord (Figure 1A, B). We generated a BAC transgenic fish line that reports the endogenous spatiotemporal expression of Pard3, a cardinal polarity protein, in live embryos (TgBAC(pard3:Pard3-EGFP)^kg301^, referred to as ‘Pard3-EGFP’). We also used an antibody for the MAGUK (membrane association and presence of the GUK domain) scaffolding molecule ZO-1 (also known as tight junction protein) on fixed tissue.

**Figure 1.**
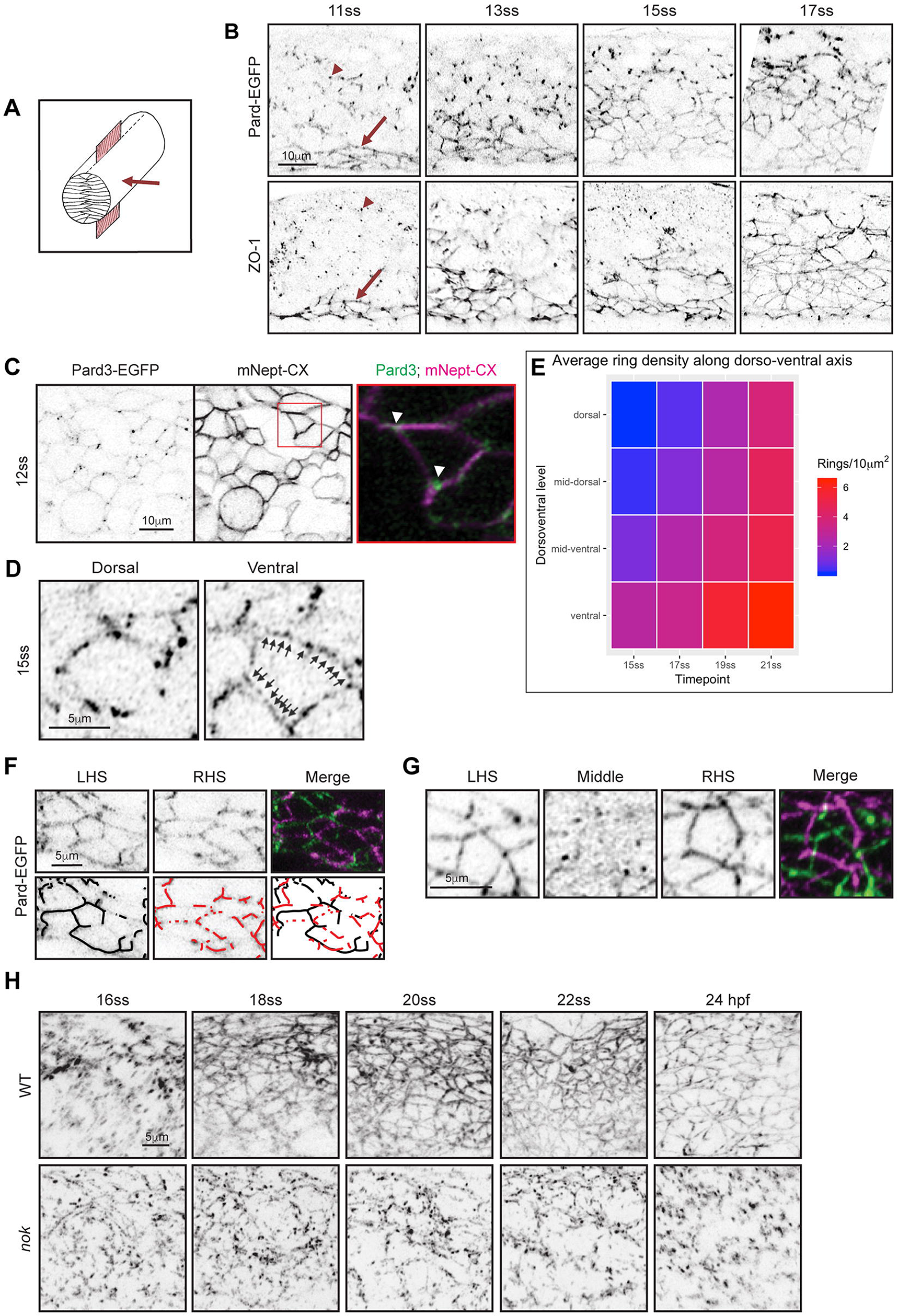
Nok dependent spot adhesion to apical ring transition. **A)** Diagram of neural rod with inserted red sheet to illustrate sagittal plane of confocal optical sections. Red arrow indicates direction of imaging. **B)** Sagittal confocal planes of neural rod in anterior spinal cord region at 11, 13, 15 and 17 somite stages. Top row images are taken from a time-lapse movie of a Pard3-EGFP embryo. Comparable images were seen at each timepoint from 3 embryos. Bottom row images are from fixed embryos stained by immunohistochemistry (IHC) for Zonula Occludens 1 (ZO-1). Comparable images were seen from 2 embryos at each timepoint. Red arrow indicates ventral apical rings and red arrowheads indicates puncta. Dorsal to the top of each panel. **C)** Sagittal confocal images at 12 somite stage showing Pard3-EGFP puncta are often located at cell vertices (arrowed, n = 8 embryos). Plasma membrane imaged using mNeptune2.5-CAAX (mNept-CX). **D** Examples of punctate Pard3-EGFP in developing apical rings from a 15 somite stage embryo. Dorsal example shows an immature, incomplete ring. Ventral example shows a more complete ring, with arrows indicating multiple puncta between vertices. Single sagittal confocal planes. **E)** A heat-map quantification of the formation of mature Pard3 apical rings from two embryos over developmental time and dorsoventral position (see methods for details of analysis). **F)** Parasagittal confocal sections from left hand side (LHS) and right-hand side (RHS) of stage 15 somite neural rod. Incomplete apical rings of Pard3-EGFP are forming independently on left and right sides of the midline. **G)** Parasagittal and sagittal confocal sections at 17 somite stage showing complete apical rings of Pard3-EGFP on either side of the midline. The sagittal section (labelled Middle) shows a largely diffuse low level of Pard3-EGFP expression in the midline territory between the left and right rings (also see supplementary movie 1). Apical ring formation illustrated in F & G was analysed from over 10 embryos. For each embryo, 10-25 apical rings were located on one side of the neuroepithelium in *en face* orientation and then a z-stack was taken through the middle of the neural rod until the rings on the opposite side were visible. We never saw an embryo containing a single plane of apical rings. There were always bilateral rings and these were always offset from each other across the midline. **H)** Images taken from confocal time-lapse movies of wild-type and *nok* morphant Pard3-EGFP embryos in sagittal orientation from the 16 somite stage to 24 hpf. Comparable images were seen from 3 embryos from each genotype and demonstrate that *nok* morphants never progress from punctate Pard3-EGFP to apical ring formation. 3 control morphants were also assessed in the Cdh2-tFT transgenic line and all had comparable apical rings to wild types.

At early neural rod stages (11 somite stage), we saw that apical rings of Pard3-EGFP and ZO-1 were first established at the midline in ventral floor plate cells (Figure 1B, e.g. arrows). In more dorsal areas of the neuroepithelium, where apical rings had not yet formed, Pard3-EGFP and ZO-1 were seen as puncta (Figure 1B, e.g. arrowheads), reminiscent of spot adherens junctions in other systems (e.g. Tepass et al., 1996). Over the next few hours of rod stage (up to 17 somite stage) the lattice work of apical rings progressively builds from ventral to dorsal such that at any given timepoint there is a developmental gradient along the dorso-ventral axis of the nascent neuroepithelium (Figure 1B, E). In addition, in contrast to a previous study (Guo et al., 2018) our results suggest that Pard3 is an early component of the nascent apical junction organisation, with both Pard3 and ZO-1 expressed in a punctate manner during a very dynamic phase of cell interdigitation across the neural keel midline (current results and Buckley et al., 2013).

### Pard3 puncta initially appear at cell vertices

To gain a better understanding of where in the cell Pard3 puncta first appear, we labelled Pard3-EGFP embryos with membrane-tagged mNeptune2.5 and imaged at neural keel stage prior to apical surface formation. We found Pard3-EGFP puncta were seen close to the midline around the perimeter of the cells, often at cell vertices (Figure 1C, e.g. arrowheads). Membrane tagged mNeptune was not enriched at these vertices, indicating the Pard3-EGFP enrichment is not simply due to more membrane at these points.

Later in development, Pard3-EGFP puncta were localised progressively closer together and not confined to cell vertices (Figure 1D). This suggests that apical rings may be built by adding Pard3 puncta at cell-cell interfaces until they coalesce to form a more or less continuous ring structure.

### *nok*^*m227*^ mutants fail to undergo puncta to apical ring transition

To begin to uncover the mechanisms that regulate formation of the apical junctions in the sagittal plane of the neural rod we examined the apical puncta to ring transition in a mutant fish line that fails to open a lumen. Nok/Mpp5a is a MAGUK scaffolding protein (Wei & Malicki, 2002), essential to retain all forms of the transmembrane protein Crumbs at the apical surface (Zou et al., 2013) and *nok* mutants have previously been reported to fail to generate an open a lumen in the neural rod and to have disorganised adherens junctions (Lowery et al., 2009, Lowery & Sive, 2005). We found that although *nok*^m227^ mutants generate Pard3-EGFP puncta in the sagittal plane at the 16 somite stage, these puncta failed to undergo the puncta-to-apical ring transition and remained as punctate deposits throughout development (Figure 1H). Nok/Mpp5a is thus critical to build apical junctional rings.

### Apical rings on either side of the tissue midline develop independently and are offset from one another

To successfully undergo *de novo* lumen formation, the zebrafish must generate not one, but two lattice-like apical surfaces in the middle of a solid tissue, one on the left side and one on the right side of the future lumen. One possibility is that a single plane of apical rings may first develop at the left-right interface and then these pioneer rings might split into left and right rings. To assess this hypothesis, we carefully analysed the development of left and right rings. Imaging sagittal and parasagittal planes we never saw an embryo containing a single plane of apical rings (Figure 1F, G, supplementary movie 1). Instead, apical rings were always found bilaterally, and incomplete apical rings were also found bilaterally on either side of the midline, showing left and right rings are built independently. Furthermore, we observed that the developing rings on either side of the tissue midline are always offset from one another, rather than having a mirror-symmetric arrangement that might be expected if individual rings split into two (Figure 1G).

### Non-sister connections impose cell offsets across the midline

Most neural rod cells derive from progenitors that divide at the midline at neural keel stages to generate mirror-symmetrically polarised sisters connected to each other across the midline (this division is called the C-division) (Buckley et al., 2013, Kimmel et al., 1994, Tawk et al., 2007). This process will at least temporarily align pairs of sister cells across the midline. However, the offset organisation of left and right apical rings later in development at the neural rod stage suggests a mechanism must exist that adds offset to this initial cell alignment. To confirm that sister cells become offset, we first generated mosaically labelled embryos and used live imaging to assess the development of the midline interface between sister cells. Soon after a C-division at the neural keel stage, sister cells appeared mirror-symmetric to each other across the tissue midline (Figure 2A, 13 somites). However, through time-lapse imaging, we observed that the shape of the nascent apical endfeet dynamically changed over time, likely in response to the movements and mitoses of neighbouring (unlabelled) cells (Figure 2A and supplementary movies 2-4). The location of the sister-cell interface in relation to the rest of the cell shape therefore also changed dynamically until the neural rod stage, when the connection between the sisters was stably offset such that the nascent apical endfeet of sister cells met at their corners (Figure 2A, 18 somites). These observations show that an initially mirror-symmetric sister cell alignment is shifted to an offset organisation following C-division mitosis. The offset conformation between C-division sister cells suggests that contralateral cell connections across the midline are likely not confined to their C-division sisters.

**Figure 2.**
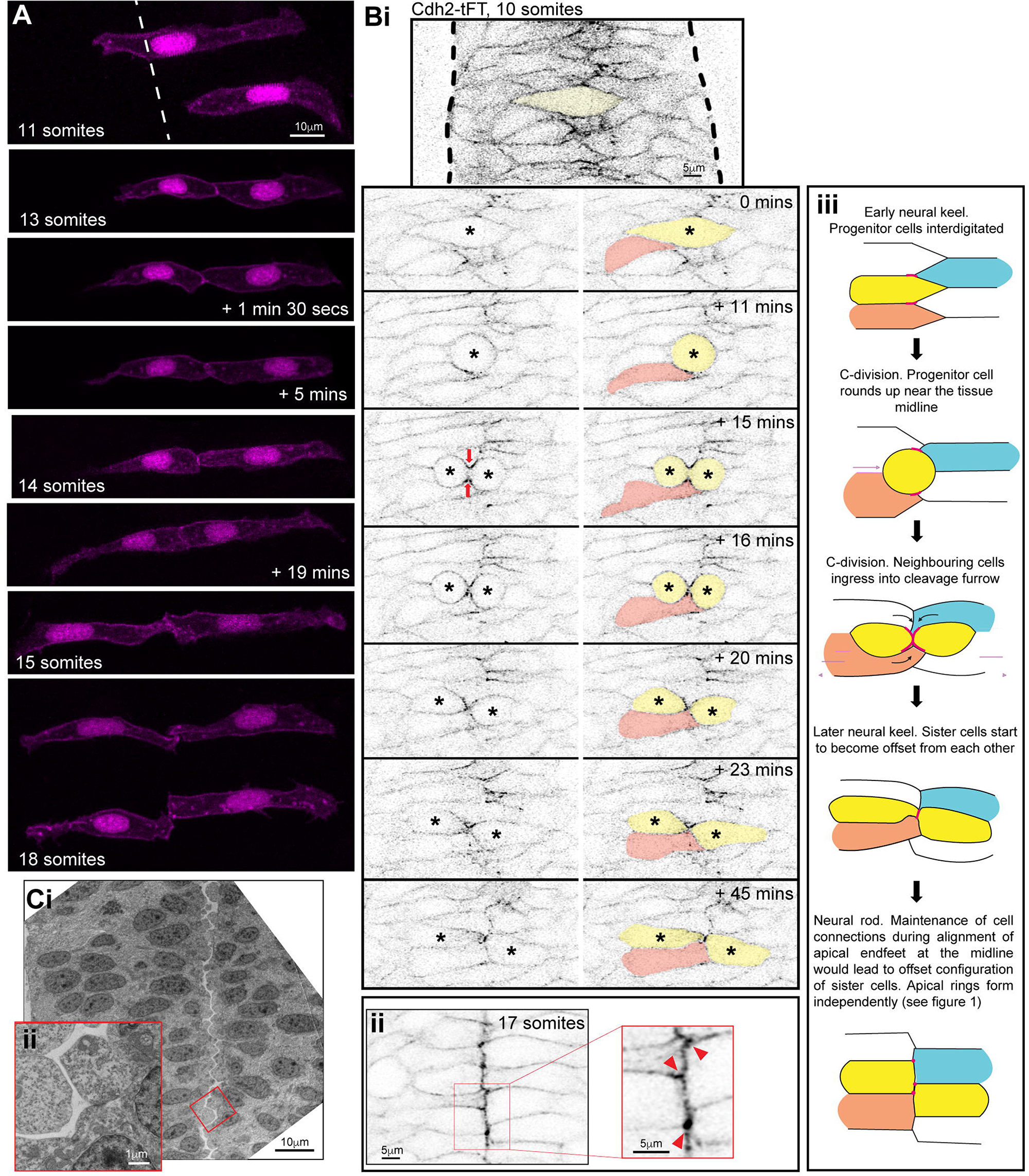
Sister-cells remain attached via their corners. **A)** Images from a time-lapse movie projection in dorsal orientation of mosaically labelled neuroepithelial cells in the hindbrain of an 11 somite stage WT embryo from neural keel stage. Membrane and nuclei are labelled in magenta. By the 13 somite stage, both cells had undergone C-division, resulting in pairs of sister cells attached across the tissue midline. The top cell pair was followed over time until the 18 somite neural rod stage, when both cell pairs were imaged. Note, overlying cells in different z-planes were artificially removed from the images to avoid obscuring the image. Overlying cells remain in the associated movies (supplementary movies 2-4). The shape of the nascent apical endfeet (and therefore the location of the connection point between sister cells in relation to the rest of the cell shape), dynamically altered over the neural keel to rod transition. However, by the neural rod stage, the connections between both pairs of cells were markedly offset and cells remained connected contralaterally via their apical corners. The configuration of cell pair connections was assessed from several different experiments at the neural rod stage (approximately 16-18 somites) and 26/31 pairs of cells from 5 embryos at neural rod stages were found to be clearly attached via their corners. The remaining 5 were either connected via a more ‘en face’ configuration or their configuration was uncertain (e.g. due to a very thin connection point). **B) i**. Single z-planes from a time-lapse movie of a C-division (yellow cell) starting at the 10 somite stage (neural keel) from a Cdh2-tFT transgenic embryo. The image contrast was increased in the reference image at the top to highlight that Cdh2 was concentrated at the interdigitation zone between cells around the tissue midline at 0 minutes. Cdh2-GFP then becomes strongly concentrated in the cleavage furrow (timepoint 15 min) and neighbouring cells ingress into the cleavage furrow (21/21 divisions, red arrows). In this example the pink cell that ingresses into the cleavage furrow gains a contralateral contact with the contralateral daughter of the C-division. Because of this contact, the contralateral daughter (yellow cell on right) becomes attached to two contralateral cells – one is its sister cell from the C-division and the other is one of its sister’s neighbouring cells (the pink cell in this example). **ii**. 5 μm projection of 17 somite stage neural rod from a Cdh2-tFT transgenic embryo. Cells are well aligned along a centrally located midline. Cdh2-GFP is upregulated along the apical midline, particularly at cell corners (red arrowheads in magnified region). **iii**. Model depicting the co-ingression of neighbouring cells into the cleavage furrow during C-division (yellow cells) and the subsequent offsetting of sister cells from each other, which precedes apical ring formation. This is based on the images similar to that shown in figure Bi. Pink lines and dots represent high levels of Cdh2 associated with the dividing cell. The ingression of either ipsilateral (e.g. orange) or contralateral (e.g. blue) neighbours into the cleavage furrow promotes the formation of multiple contralateral connections across the midline. For example, the left-hand yellow sister cell becomes attached to both its right-hand yellow sister cell and the ingressing blue cell. The right-hand yellow sister cell becomes attached to both its left-hand yellow sister cell and the ingressing orange cell. This is one way in which multiple ipsilateral and contralateral adhesions might be promoted during C-division mitosis **C) i**. Transmission electron micrographs of a 20 hpf embryo hindbrain in transverse orientation. The lumen has just started to open from the midline. The interface between contralateral cells has a striking ‘zig-zag’ pattern (3/3 19-20 hpf embryos). **ii**. Inset magnified region from **i**.

To examine the dynamics of ipsilateral and contralateral cell adhesions during the process of the midline C-division, we took advantage of a BAC transgenic line expressing the cell adhesion protein Cdh2 fused to a tandem fluorescent timer (Revenu et al., 2014) and we imaged Cdh2-GFP from this line. Cdh2-GFP is expressed in all neural rod cells over the entire plasma membrane. Prior to C-divisions, cells intercalated across the rod midline and Cdh2-GFP was enriched in this interdigitation zone, showing that cells contact and adhere to several contralateral, as well as ipsilateral, cells (Figure 2Bi). When cells entered C-division mitosis at or close to the middle of the neural keel, we found that cell processes from neighbouring cells ingressed into the cleavage furrow and distinct enrichments of Cdh2-GFP became largely localised to the cleavage furrow between sister cells during telophase (red arrows, Figure 2Bi). This cell ingression allowed neighbouring cells to contact and presumably adhere to both the ipsilateral and contralateral daughter of the C-division. Cell ingression into the cleavage furrow therefore provides a potential mechanism to promote connections between multiple contralateral and ipsilateral cells (as cartooned in figure 2Biii).

By the 17 somite stage, Cdh2-GFP is enriched at the interface between left and right cells, sometimes at the apical cell vertices (Figure 2Bii). By this stage the left-right interface is now largely a straight line at the midline of the neural rod. To see whether the offset cell conformation between cells on the left and right of the neural rod persists until lumen opening, we looked at electron micrographs taken at the very start of lumen inflation. This showed a striking zig-zag conformation between cells separating across the apical midline (Figure 2C). It therefore appears that the offset conformation between left and right cells persists until the lumen inflates, separating the left and right sides of the neural rod.

In summary, this data suggests that the C-division can promote connections between multiple ipsilateral and contralateral cells during epithelialisation. We suggest that the maintenance of these connections until rod stage would promote an offset conformation between sister cells.

### A single midline interface enriched in polarity and cell adhesion proteins precedes the formation of bilateral apical rings

To better understand the transition from spot-like junctional specialisations to two independent lattices of apical rings on left and right sides of the neural rod midline we analysed the changing distribution of Pard3 and the adhesion protein Cdh2 during this transition. Since there is a developmental gradient along the dorso-ventral axis of the neuroepithelium (Figure 1B and E), we focused on a mid-dorsoventral plane of the anterior spinal cord. We imaged Pard3-EGFP and Cdh2-GFP from neural keel through to neural rod stages in the horizontal plane (Figure 3Ai-iii). This showed that Pard3-EGFP and Cdh2-GFP initially have different temporal patterns of expression. At the 11 somite stage (14 hpf, neural keel stage) puncta of Pard3-EGFP were localised broadly around the tissue midline, along the interfaces of interdigitating cells. These correspond to the spot distribution of Pard3-EGFP seen in sagittal sections (Figure 1B and C). At the same timepoint Cdh2-GFP was expressed at low level throughout the cell membrane but also slightly elevated in the interdigitation zone, mostly in discrete spots. By the 17 somite stage (17.5 hpf, neural rod stage), both Cdh2-GFP and Pard3-EGFP were clearly enriched at the midline in a near continuous, fairly straight expression domain at the interface between left and right cells. Consistent with a developmental gradient along the dorso-ventral axis of the neuroepithelium, dorsal to this plane, Pard3-EGFP was still expressed in a discontinuous mediolateral stripe pattern, and ventral to this plane it appeared in two parasagittal domains either side of the tissue midline (Figure 3Av). The intermediate level that shows a single left-right interface of Pard3-EGFP expression occupies only a small and transient dorsoventral domain that progresses dorsally as the assembly of bilateral rings progresses from ventral to dorsal.

**Figure 3.**
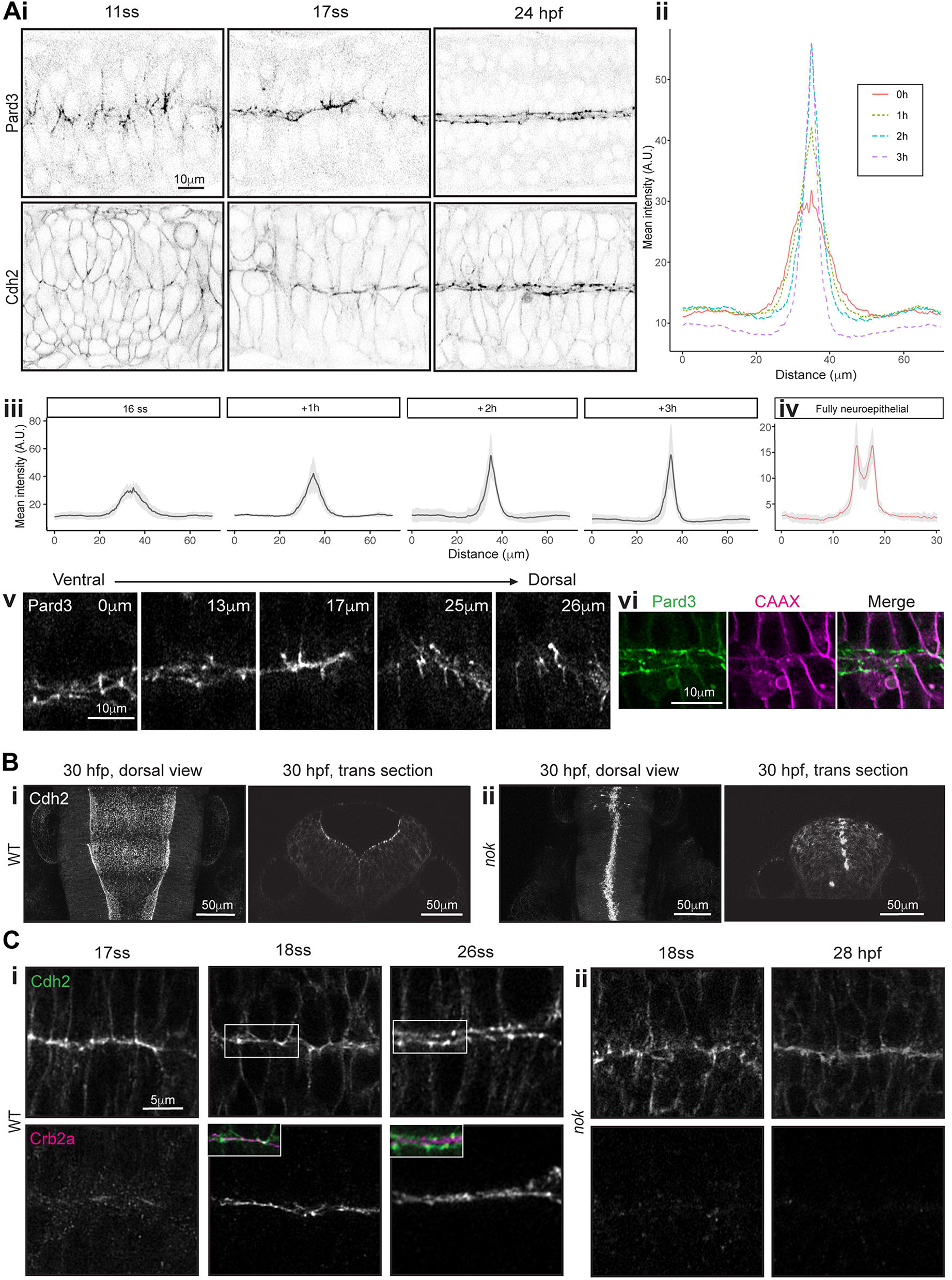
Nok dependent remodelling of midline adhesions. **A) i**. Images taken from confocal time-lapse movies of Pard3-EGFP and Cdh2-GFP embryos in horizontal orientation at 11 somite, 17 somite and 24 hpf stages. Comparable images were seen from 3 embryos from each transgenic line. **ii** and **iii**. Mean intensity profiles from 6 embryos, quantifying Pard3 intensity across the basal-to-basal width of the developing neuroepithelium over time, starting at the 16 somite stage. Standard deviation is shown as a grey ribbon around the line profile for each timepoint in **iii**. See methods for details of analysis. The midline expression domain of Pard3 increases in intensity and decreases in width over development. **iv**. Mean intensity profiles from the same 6 embryos, quantifying Pard3 intensity across the basal-to-basal width of the neuroepithelium at the fully neuroepithelial stage. A parallel ‘tramline’ of Pard3-EGFP expression shows the two apical sheets on each side of the neural tube. **v**. Horizontal confocal planes of 17 somite stage neural rod showing Pard3-EGFP expression at five different dorsoventral levels. The single elevated plane of expression at the left right interface seen at level 17 μm, lies dorsal to levels where apical rings are already formed and ventral to levels where expression is more prominent in mediolateral streaks. **vi**. Single horizontal plane confocal section of Pard3-EGFP and mNeptune2.5-CAAX and merge at approximately 24 hpf. Plasma membranes meet at the tissue midline while Pard3 is now largely located in two parallel parasagittal domains. **B)** Horizontal and transverse confocal sections of 30 hpf. **i**. wild-type and **ii**. *nok*^*m227*^ mutant Cdh2-GFP embryos at the hindbrain level. The hindbrain lumen remained closed in 5/5 *nok*^*m227*^ mutant embryos analysed and is always open in wild type embryos. **C)** Horizontal confocal sections of **i**. wild-type and **ii**. *nok*^*m227*^ mutant Cdh2-GFP embryos at the anterior spinal cord level, stained by IHC for Crb2a. Insets in the 18 and 26 somite stages of wildtypes show a merge of Crb2a and Cdh2-GFP expression. **i**. In wild-type embryos, Cdh2 and Crb2a were initially colocalised at the tissue midline at the 18 somite stage (4/4 embryos) but Cdh2-GFP protein was later displaced basolaterally to form two independent stripes of expression either side of the midline by the 26 somite stage (8/8 embryos). In *nok*^*m227*^ mutants, Crb2a was not present at the midline at the 18 somite stage (4/4 embryos) and Cdh2-GFP remained in a single expression domain at the tissue midline even as late as 28 hpf (5/5 embryos).

To analyse the distribution of cell membranes relative to the expression of Pard3-EGFP, we revealed membranes by injecting TgBAC(pard3:Pard3-EGFP)^kg301^ embryos with mRNA mNeptune2.5-CAAX at the 1-2 cell stage. By 24 hpf, although spinal cord cells at almost all dorsoventral levels do still meet at the left-right interface (see membrane label in Figure 3vi), Pard3-EGFP and Cdh2-GFP no longer localise to this interface; rather their expression has fully transitioned from mediolaterally arranged puncta through a single midline expression domain and into two parasagittal domains either side of the tissue midline (Figure 3Ai, iv). These two domains are characterised by periodically elevated spots of expression that correspond to points within the left and right apical rings. Much lower levels of both Pard3-EGFP and Cdh2-GFP remain in the medial zone between the bilateral apical rings (Figure 3Ai, iv).

These data suggest left and right cells initially adhere together across the midline through Cdh2-rich adhesions, but this interface is then largely cleared of both Cdh2 and Pard3 as these adhesion and polarity proteins are redeployed more basally to the apicolateral border where they contribute to adhesive junctions between ipsilateral cells. This basolateral redeployment of junctional proteins is likely to be a critical step in the process of releasing cells from their contralateral adhesions that allows lumen formation.

### Nok/Mpp5a is required to redeploy adhesions to the apicolateral border

Work in *Drosophila* epithelia has shown that Crumbs is required for the proper localisation of Baz (*Drosophila* Pard3) and E-cadherin at apical junctions, displacing them from the apical-most surface of single sheet epithelial cells to the apico-lateral border of the cell (Grawe et al., 1996, Harris & Peifer, 2005, Morais-de-Sa et al., 2010, Tepass, 1996). Although eliminating Crumbs in zebrafish is difficult because there are multiple Crumbs proteins, we took advantage of the *nok*^*m227*^ mutant that is unable to localise Crumbs proteins to the apical surface (Zou et al., 2013). Since we earlier demonstrated that the apical spot to ring transition did not occur in *nok*^*m227*^ mutants (Figure 1H), we next wanted to test the role for Nok in displacing Pard3 and Cdh2 proteins from a single line at the left-right interface, to the two bilateral expression domains seen at more mature time points. To do this we generated a *nok*^*m227*^ mutant; Cdh2-tFT compound transgenic line. Whereas ‘wild-type’ Cdh2-tFT embryos opened a lumen in hindbrain and anterior spinal cord (Figure 3Bi), *nok*^*m227*^; Cdh2-tFT embryos failed to open a lumen (Figure 3Bii), in line with previous observations on *nok* mutant embryos (Lowery & Sive, 2005).

To understand the *nok*^*m227*^ phenotype in relation to Cdh2 remodelling and its relation to Crumbs expression, we examined the localisation of both proteins in wild-type and *nok*^*m227*^ embryos. In wild-type embryos (Figure 3Ci) Crumbs2a is first expressed at the neural rod 17/18 somite stage at the left-right interface, coincident with a single line of expression of Cdh2. However, by the 26 somite stage, when Crumbs2a protein was still localised to the left-right interface, Cdh2-GFP protein was displaced to form two slightly more basolateral lines of expression. The Cdh2-GFP expression was located immediately lateral to Crumbs2a, with very little overlap of expression.

In *nok*^*m227*^ mutant embryos (Figure 3Cii) Crumbs2a expression is absent from the midline as expected. At the neural rod 18 somite stage *nok*^*m227*^ mutant Cdh2-tFT embryos showed a single domain of enrichment of Cdh2-GFP at the left-right interface similar to wild-type Cdh2-GFP embryos. However, unlike wild-type embryos, Cdh2-GFP expression remained as a single midline domain in *nok*^*m227*^ mutants and failed to undergo the transition to two lines of Cdh2-GFP either side of the tissue midline, even up to stages as late at 28 hpf.

Together, these data show that left and right cells initially adhere together across the midline through contralateral adhesions. However, both Cdh2 and Pard3 adhesion and polarity proteins are then redeployed more basally to the apicolateral border where they contribute to adhesive junctions between ipsilateral cells to build coherent, bilateral sheets of neuroepithelial cells. This basolateral redeployment of junctional proteins depends on Nok/Mpp5a and is likely to be a critical step in the process of releasing cells from their contralateral adhesions to allow lumen formation. We propose this function of Nok/Mpp5a is likely to operate through its ability to localise Crumbs proteins to the left-right interface. This is consistent with the loss of Crumbs proteins at the left-right interface and the loss of basal displacement of junctional proteins away from the left-right interface in *nok*^*m227*^ mutants (Figure 3C). This allows the persistent adhesion between cells on left and right sides, which is likely to cause the inhibition of lumen formation in *nok*^*m227*^ mutants.

### Persistent adhesions may contribute to lack of epithelial maintenance in Nok and Rab11a deficient epithelia

To test the hypothesis that Nok deficient cells retain persistent adhesions across the midline, we analysed the behaviour of clonally related cells in the hindbrain regions of 25 hpf wild-type embryos that had open lumens and *nok* morphant embryos that were unable to open a lumen. In wild type embryos, cells had successfully separated across the midline following C-division; they maintained an elongated morphology that stretched from apical to basal surface over the course of imaging and later differentiative divisions (D-divisions) occurred at the apical (ventricular) surface (Figure 4A). However, in *nok* morphant tissue, cells were not separated across the midline, often did not fully extend to the basal surface and they appeared clumped together, often in rosette-like structures. D-divisions in *nok* morphants occurred within these cell clumps, often at a distance from the midline of the tissue, in contrast to wild-type cells (Figure 4B and quantified in 4C). The cell behaviour of ectopic divisions in *nok* morphants is accompanied by a lateral dispersal of Pard3 expression from the midline (Figure 4D). Thus, as well as being unable to undergo the puncta-to-apical ring transition (Figure 1H), Nok deficient cells show persistent adhesions and apical midline organisation becomes progressively fragmented over time.

**Figure 4.**
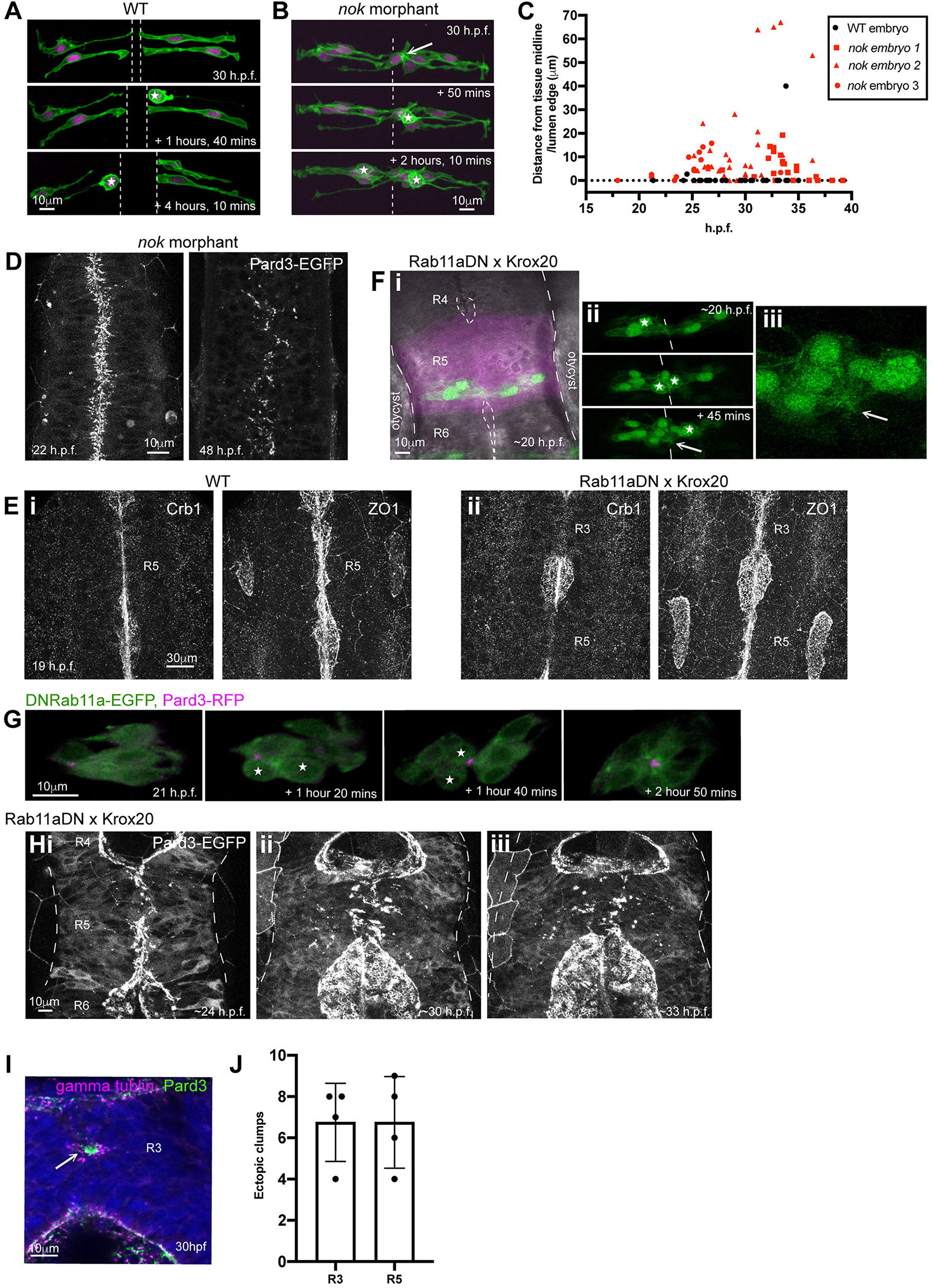
Persistent adhesion in Nok and Rab11a deficient cells. **A)** Images from a time-lapse movie of mosaically labelled neuroepithelial cells in the hindbrain of a 30 hpf WT embryo in dorsal orientation. Cells have separated across the tissue midline and the lumen has opened (short dotted lines). D-divisions (stars) occur at the apical surface and cells re-established an elongated morphology towards the basal surface following division. **B)** Images from a time-lapse movie of mosaically labelled neuroepithelial cells in the hindbrain of a 30 hpf *nok* morphant embryo in dorsal orientation. Cells have failed to separate across the tissue midline (dotted line) and formed clumps (sometimes with a rosette-like structure, arrow. D-divisions (stars) occurred near the centre of the cell clumps, which was often not situated near the tissue midline. Of the 117 daughter cells that we were able to follow post division, at least 40% did not fully re-extend to the basal side of the neural rod. **C)** Graphical representation of individual cell division locations in relation to the tissue midline or lumen edge over development. 71 cells were analysed from 3 *nok* morphant embryos. In embryos over 25 hpf, 49% of *nok* morphant cells divided 5μm or more away from the tissue midline. Division locations from a single wild type embryo example have been included in black for reference. However, wild type divisions always occur exactly at the tissue midline or lumen surface, except for rare basal divisions (McIntosh et al., 2017). **D)** 10 μm z-projection in dorsal orientation at a mid dorso-ventral level through the hindbrain of *nok* morphant Pard3-EGFP embryos at 22 hpf and 48 hpf. At 22 hpf Pard3-EGFP was localised near the tissue midline but did not form continuous straight expression domains as seen in wild types (Figure 3Ai) and the lumen failed to open. By 48 hpf Pard3-EGFP localisation became fragmented into clumps. 9/9 *nok* mutant embryos and 6/8 *nok* morphant embryos over 24 hpf old from 5 different experiments had fragmented midlines and ectopic apical proteins. The extent of this disorganisation was greater in older embryos. 2/8 *nok* morphant embryos had a milder phenotype, with a few lumen openings present and no overtly ectopic apical proteins (see supplementary figure 1). 9/9 wild type embryos had normal apical surfaces with no ectopic apical proteins. **E)** 70-80 μm z-projections in dorsal orientation in the hindbrain of 19 hpf embryos, stained via IHC for Crumbs 1 (Crb1) and ZO1. **i**. Wild-type embryo. Both Crb1 and ZO1 were localised to the apical midline (n=3/3). **ii**. Embryo in which UAS:DNRab11a was expressed under the Krox20:KalTA4 activator in rhombomeres 3 and 5 (DNRab11a x Krox20). ZO1 was localised to the midline but Crb1 was largely absent from rhombomeres 3 and 5 (n=3/3). **F) i**. 11 μm z-projection in dorsal orientation of mosaically labelled neuroepithelial cells in the hindbrain of a 20 hpf DNRab11a x Krox20 embryo. Single z-section in greyscale and 52μm projection in magenta denoting DNRab11a rhombomere 5 for reference. Long dotted lines denote basal surfaces. The lumen in rhombomere 5 failed to open but had started to open in wild-type rhombomeres 4 and 6 (short dotted lines). Note, overlying cells in different z-planes were artificially removed from the images to avoid obscuring the image. **ii**. Images from a time-lapse movie of the cells in i, which had failed to separate across the tissue midline (midline is indicated by white dashes). Similar to *nok* morphant embryos (Figure 4B), following D-divisions (stars), cells remained attached and were arranged into a rosette-like structure (arrow), enlarged in **iii**. Cell clumping or rosette-like structures were observed in the DNRab11a rhombomeres of all 9 embryos analysed from 3 different experiments. **G)** Time-lapse 3D reconstruction showing an example of mosaically distributed DNRab11a-EGFP neuroepithelial cells in the hindbrain of a 24 somite stage (equivalent to 21 hpf) embryo. Following D-divisions (stars), cells did not separate and formed a rosette around a centrally located puncta of Pard3-RFP. **H)** Approximately 70 μm z-projections from a time-lapse movie through a DNRab11a x Krox20 embryo hindbrain labelled with Pard3-EGFP fusion protein, starting at approximately 24 hpf. In the same way as in *nok* morphants in figure 4D and as previously reported (Buckley et al., 2013), Pard3 was initially localised near the tissue midline of rhombomere 5, but the lumen failed to open and Pard3 localisation became fragmented into clumps that were localised progressively further from the tissue midline. Whilst the majority of the DNRab11a x Krox20 embryos that we tested from 4 different experiments had fully penetrant phenotypes, even in heterozygotes (17/17 had fully closed lumens and ectopic apical proteins in DNRab11a rhombomeres 3 and 5), 9/10 embryos from 2 of our later experiments did not have a fully penetrant DNRab11a phenotype. In these embryos some lumen opening was present, demonstrating that cells were able to separate from each other across the midline. In this case no or very little ectopic apical protein was seen. We think that this lower DNRab11a penetration was due to methylation of the Tg(UAS:mCherry-Rab11a S25N)^mw35^ line since, when we bred previous generations of the line, we saw fully penetrant phenotypes again. This correlation between extent of DNRab11a penetrance, lumen opening and ectopic apical proteins was also seen in *nok* morphant embryos (supplementary figure 1). **I)** 4 μm z-projection in dorsal orientation 23 μm below the rhombomere surface of a 30 hpf DNRab11a x Krox20 embryo hindbrain at rhombomere 3, stained via IHC for Pard3 and gamma-tubulin. Pard3 is localised in a round clump, surrounded on all sides by centrosomes. **J)** Both DNRab11a rhombomeres 3 and 5 had approximately 7 ectopic clumps of Pard3 and gamma-tubulin at 30 hpf (n=4 embryos. Error bars denote standard deviations). We did not see ectopic clumps in wild-type rhombomeres.

Since we hypothesise that Nok functions through its ability to localise Crumbs to the nascent apical surface, we next analysed cell behaviours following another manipulation that depletes Crumbs from the apical surface. For this we expressed a dominant-negative (DN) form of Rab11a in rhombomeres 3 and 5. We previously showed that this manipulation reduces apical Crumbs 2a localisation and prevents lumen opening in these segments (Buckley et al., 2013). Here, we show that apical Crumbs 1 is also downregulated in DNRab11a rhombomeres (Figure 4E), confirming that this manipulation is sufficient to downregulate apical trafficking of multiple Crumbs paralogs. We found that cells in the DNRab11a rhombomeres did not separate following D-division and formed cell clumps and rosette-like structures (Figure 4F). An example of rosette formation around a central focus of Pard3 is depicted in Figure 4G. Similar to *nok* morphants (Figure 4D), at the tissue level, a lateral dispersal of Pard3-EGFP from the midline was seen, progressively fragmenting the midline organisation over time (Figure 4H). This is in support of our previous observations of ZO-1 dispersion in DNRab11a segments (Buckley et al., 2013). Immunohistochemical (IHC) staining of DNRab11a rhombomeres revealed ectopic clumps of Pard3 and centrosomes, often arranged in rosette-like structures with centrally located Pard3 surrounded by centrosomes (Figure 4I and J).

Interestingly, whilst cells situated in the middle of closed-lumen DNRab11a rhombomeres 3 and 5 formed clumps, cells at the edges of these rhombomeres often had a typical elongated neuroepithelial morphology and were able to build an aPKC positive apical surface that was contiguous with the wild-type apical surface in adjacent open-lumen wild-type rhombomeres 2, 4 and 6 (Figure 5A). Edge cells were oriented obliquely along the anterior-posterior axis of the embryo, apical towards the open lumens (e.g. arrows in figure 5Aii). To determine how this morphology arose, we made time-lapse movies of DNRab11a/wild-type interface neuroepithelial cells. We found that interface cells remained connected across the midline as the neighbouring wild-type lumen inflated, acting like a hinge and reorienting the cells at the boundary between DNRab11a and wild-type rhombomeres towards the opening lumen (Figure 5Bi). Despite these persistent connections between cells, the DNRab11a luminal surface expanded via cell division and subsequent reintegration into the epithelium (Figure 5Bii). Cell divisions in the centre of DNRab11a rhombomeres were disorganised and did not orient along the midline of the tissue, whilst those occurring at the DNRab11a luminal surface occurred with parallel orientation Figure 5Biii.) These results demonstrate that DNRab11a cells at the DNRab11a/wild-type interface are able to successfully reintegrate into the epithelium following division if they have access to an apical lumen surface.

**Figure 5.**
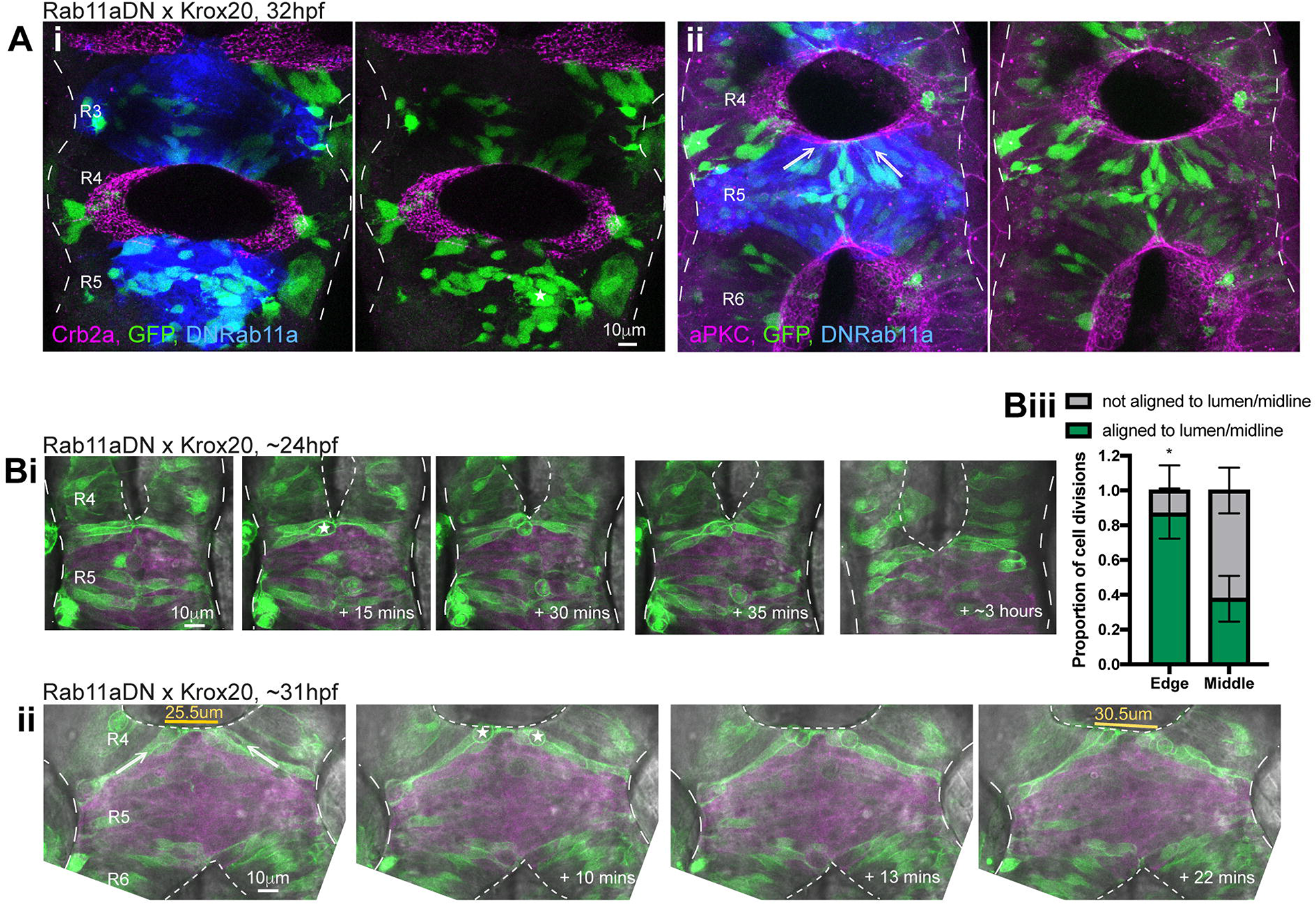
Cells at the WT/DNRab11a interface contribute to the luminal surface. **A)** 15-18 μm z-projections through 32 hpf DNRab11a x krox20 embryo hindbrains stained by IHC for Crb2a (i) or aPKC (ii). Cells are mosaically labelled with cytoplasmic GFP and H2A-GFP. **i**. Crb2a is largely (but not completely) absent from DNRab11a apical regions (5/5 embryos). **ii**. aPKC is present around the whole apical surface of both wild-type and DNRab11a rhombomeres (4/4 embryos). Cells situated in the centre of DNRab11a rhombomeres clumped together (e.g. star in i) whilst cells in contact with the open lumens had elongated morphology (e.g. arrows in ii). **B)** Single z-slices from a time-lapse movie in dorsal orientation of mosaically labelled neuroepithelial cells in rhombomere 5 of an approximately 24 hpf DNRab11a x Krox20 embryo over time. Long dotted lines denote basal surfaces. Short dotted lines denote apical surfaces. **i)** As the neighbouring lumen inflated, cells at the edge of the DNRab11a rhombomere remained connected across the midline and cells near the edge divided at this central point of connection with parallel orientation (star). **ii)** As the neighbouring lumen inflated further, cells near the WT/DNRab11a interface reoriented towards the open lumen (e.g. arrows). Cells divided at the opening luminal surface with parallel orientation (stars). This widened the DNRab11a luminal surface further (see measurements in orange). **iii)** Quantification of cell division orientation. 104 DNRab11a cell divisions were analysed from 3 embryos over approximately the 24-40 hpf period of development. 87% of DNRab11a cells dividing at the luminal edge did so parallel to the opening lumen, whilst only 38% of DNRab11a cells dividing in the middle of rhombomeres 3 and 5 did so parallel to the midline (P=0.0121, unpaired, 2-tailed t-test). Error bars denote standard deviations between embryos.

Our data suggests that persistent adhesions between cells in Nok deficient and DNRab11a embryos both inhibits lumen formation and promotes the formation of local clumps of cells. We earlier showed that adhesions between cells in Nok deficient and DNRab11a embryos are not reorganised into a lattice of ring-like adhesions between ipsilateral junctional belts (Figure 1H), whose role is to generate and maintain a continuous sheet of cells. We found that many cells undergo ectopic divisions in this environment, presumably further contributing to ectopic cell clump formation and loss of epithelial structure (Figure 4B and C). However, despite persistent adhesions between DNRab11a cells they could integrate into a neuroepithelium if they had access to an apical lumen surface initiated by wild-type cells (Figure 5). Therefore, together, our data suggests that a combination of a lack of cell separation following division and a lack of apical ring formation results in the progressive fragmentation of midline organisation over successive rounds of division in Nok deficient and DNRab11a embryos.

### Nok is necessary for apical ring formation even in the absence of cell division

A recent paper’s analysis of cell divisions and lumen formation in *nok*^*m520*^ mutants suggested that Nok/Crumbs function in lumen formation may be specifically related to resolving connections between sister cells following the C-division (Guo et al., 2018). In the course of analysing junction formation in *nok*^*m227*^ mutants, we uncovered that the loss of lumen formation itself in these mutants was more complicated than expected. In line with previous observations (Lowery et al., 2009, Lowery & Sive, 2005), we found the loss of lumen formation was not fully penetrant and consistently found that small lumens formed at the level of the midbrain hindbrain boundary (Figure 6A). Less frequently small lumens also formed in the dorsal aspects of the hindbrain at the level of the otic vesicles (Figure 6B). We found that Crumbs 2a protein was present at the apical surface of these lumens (Figure 6A), consistent with the role of Crumbs in the successful formation of an apical surface and opening of a lumen.

**Figure 6.**
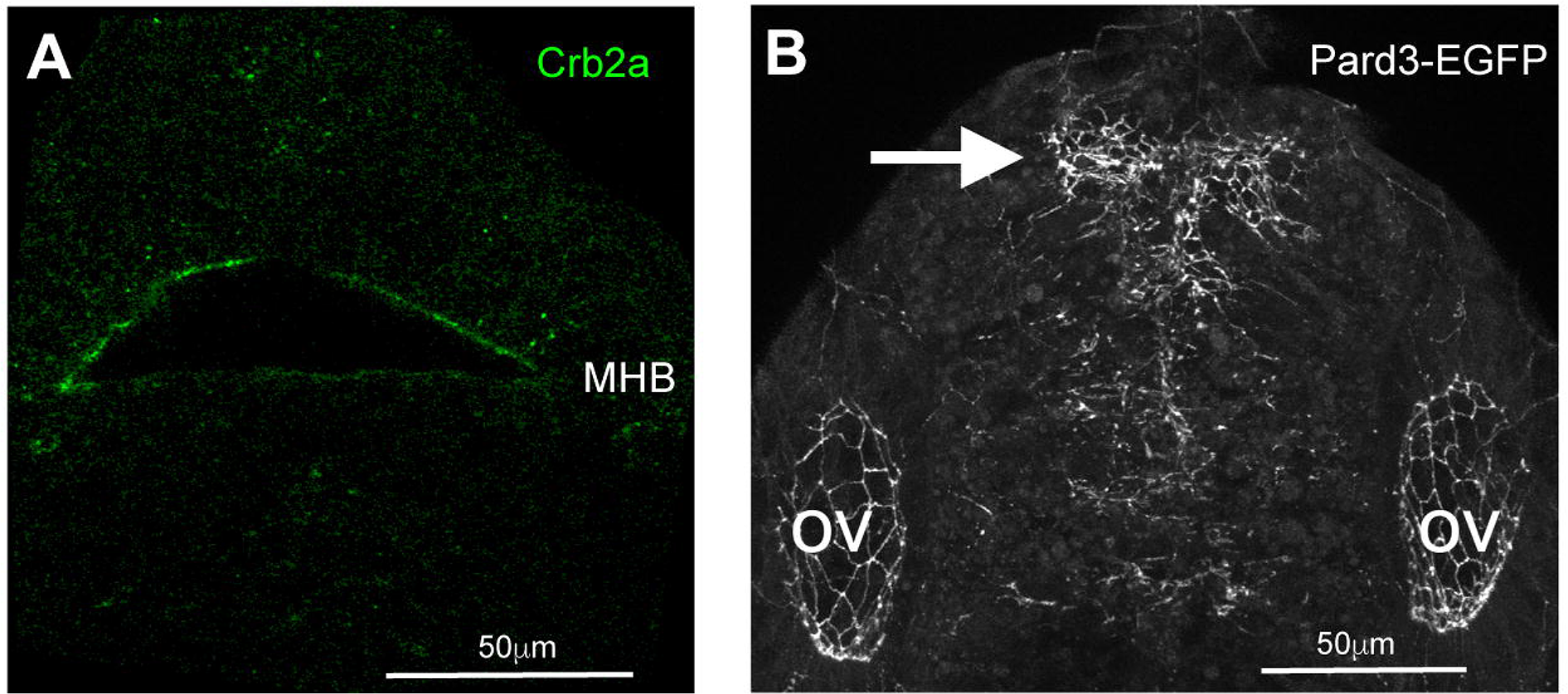
*nok*^*m227*^ mutants still open a lumen at the midbrain-hindbrain boundary. A. Single horizontal z-plane of the hindbrain neural rod from a *nok*^*m227*^ mutant embryo at the level of the midbrain-hindbrain boundary (MHB), imaged for Crb2a immunoreactivity. A lumen has opened, which is lined with Crumbs2a protein apically (5/5 embryos). B. A 57 μm maximum projection of a z-stack of the hindbrain neural rod from a 28 somite stage *nok*^*m227*^ mutant; Pard3-EGFP compound transgenic embryo at the level of the otocysts. Apical rings have formed at the lumen surface (arrow, 4/4 embryos).

Since the closed-lumen phenotype is not fully penetrant in *nok* mutants, we suggest that using the rescue of the open lumen phenotype is problematic in *nok*. To better analyse whether Nok has specific functions in relation to the C-division we therefore analysed apical ring formation in the anterior spinal cord. This region is distant from the midbrain hindbrain boundary and the level of otic vesicles and provides a more reliable and precise assay of Nok function. We have shown that Nok and Rab11a are required for cells to separate across the midline following the C-division. We have previously demonstrated that left-right separation was not rescued by blocking division in DNRab11a embryos, suggesting that the role of Rab11a is not exclusively related to cell-cell separation between sister cells (Buckley et al., 2013). To address whether Nok/Crumbs function may be specifically related to resolving connections between sister cells (as suggested in Guo et al. 2018), we analysed the key events of apical ring formation in *nok*^*m227*^ mutant embryos with and without C-divisions. We blocked C-divisions using the S-phase inhibitor Aphidicolin and analysed Pard3-GFP expression in the anterior spinal cord region. Apical rings of Pard3-EGFP are not rescued by blocking C-divisions in *nok*^*m227*^ mutants (Figure 7). The efficacy of the division block was confirmed by the enlarged nuclear size in aphidicolin treated embryos (Figure 7F and G) and the fewer, enlarged apical rings in the aphidicolin treated siblings (Figure 7B and H). These results demonstrate that Nok function is not specifically related to remodelling adhesions between C-division sisters and show that Nok plays a more fundamental role in the formation of apical junctional rings and release of contralateral adhesions during *de novo* polarisation of an epithelial tube.

**Figure 7.**
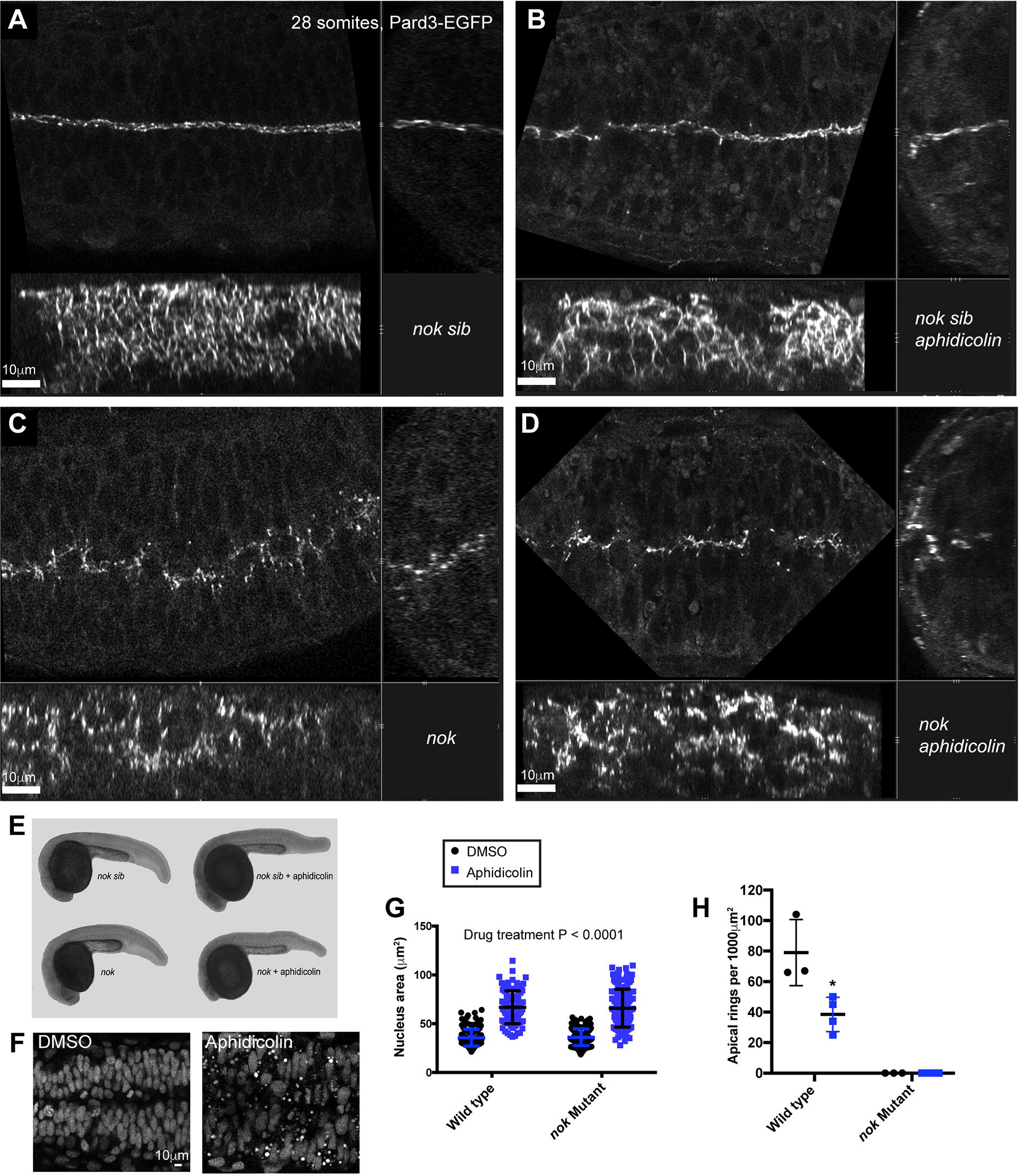
Nok is necessary for apical ring formation even in absence of cell division. All embryos are at 28 somite stage. **A-D)** Orthogonal series of horizontal (top left), transverse (top right) and sagittal (bottom) confocal planes of Pard-EGFP expression in the following embryos: **A)** wildtype sibling treated with DMSO vehicle control. Many small apical rings have formed at the tissue midline (n=3/3 embryos, quantification in F). **B)** wildtype sibling treated with aphidicolin to block the C-divisions. Fewer, large apical rings have formed at the tissue midline (n=4/4 embryos, quantification in F). **C)** *nok*^*m227*^ mutant embryo treated with DMSO vehicle control. No apical rings have formed at the tissue midline. Pard3 is visible as spots along the midline plane (n=3/3 embryos, quantification in F). **D)** *nok*^*m227*^ mutant embryo treated with aphidicolin to block C-divisions. No apical rings have formed at the tissue midline. Pard3 is visible as spots along the midline plane (n=4/4 embryos, quantification in F). **E)** Overall body shape of wildtype siblings and *nok*^*m227*^ mutants with and without aphidicolin treatment. **F)** Horizontal section through neural tube showing nuclear staining in wildtype embryos treated with DMSO or aphidicolin to block C-divisions. Aphidicolin-treated embryo have larger nuclei, demonstrating that S-phase division-block was successful (quantification in G). **G)** Quantification of nucleus area of wildtype siblings and *nok*^*m227*^ mutants with and without aphidicolin treatment. Data from 100-150 nuclei was pooled from 2-3 embryos in each group and analysed by 2-way ANOVA. Areas of Aphidicolin treated nuclei were, on average, 30 μm^2^ bigger than DMSO treated embryos (P<0.0001). There was no significant difference in nuclei area between wild type and *nok*^*m227*^ mutant embryos (P=0.8561). Error bars denote standard deviations. **H)** Quantification of apical ring number in wildtype siblings and *nok*^*m227*^ mutants with and without aphidicolin treatment. Numbers of apical rings per 1000 μm^2^ were calculated from 3-4 embryos per group. No apical rings were seen in any of the *nok*^*m227*^ mutants. There were, on average, 40 fewer apical rings per 1000 μm^2^ in aphidicolin treated wild type embryos than in DMSO treated wild type embryos (unpaired t-test, P=0.0222). Error bars denote standard deviations.

## Discussion

Our work uncovers three aspects of *de novo* apical surface generation within a complex organ *in vivo*. First despite the very different cellular organisations, apical surface generation at the centre of a solid organ primordium follows a strikingly similar sequence of spot to ring junction building to those previously described for apical surface generation at a free surface of a sheet of cells. Second the displacement of proteins from the nascent apical surface to the nascent apicolateral junctional belt domain serves two purposes in a solid organ primordium; it simultaneously contributes to the building of epithelial junctional belts and releases adhesions with contralateral cells that would otherwise inhibit the generation of a free surface and lumen formation. Our comparable results in *nok* ^*m227*^ mutant/morphant and DNRab11a tissue lead us to propose that the correct apical localisation of Crumbs protein is necessary for both apical ring formation and the resolution of adhesions between cells, defects in which result in the progressive fragmentation of the apical midline over the course of multiple rounds of cell division. Thirdly neuroepithelial progenitor cell divisions across the nascent apical plane (C-divisions) do not promote exclusive connections between contralateral sister cells. Instead connections between multiple ipsilateral and contralateral partners are promoted by neighbour ingression into the cleavage furrow of the C-division and appear to contribute to the offset alignment of contralateral cells across the midline. Through this realignment, sister cells do not remain connected ‘*en face*’ as previously suggested (Guo et al., 2018), but instead sisters become attached in an offset configuration from one another across the midline. In addition, we address a recent suggestion that the MAGUK scaffolding protein Nok is only required to remodel apical protein location following cell divisions across the nascent apical plane (Guo et al., 2018). We show this is not the case as Nok-dependent apical protein remodelling is required for apical surface generation both with and without cell divisions.

### The roles of Nok and Rab11a

During epithelial development of the *Drosophila* ectoderm, Crumbs is required for the proper localisation of Baz (*Drosophila* Pard3) and E-cadherin at apical junctions, displacing them from the apical-most surface of single sheet epithelial cells to the apico-lateral border of the cell (Grawe et al., 1996, Harris & Peifer, 2005, McGill et al., 2009, Morais-de-Sa et al., 2010, Pilot et al., 2006, Tepass, 1996) and thus building the apicolateral junctional belts that tie the epithelial cells together (St Johnston & Ahringer, 2010, Wei et al., 2005).

Here, we describe a similar displacement of Pard3 and repositioning of the main zebrafish neuroepithelial cadherin, Cdh2, to the apicolateral cell borders in the zebrafish neural rod. In this case the displacement occurs at the midline plane within a solid rod of cells rather than at a free surface, but despite this fundamental difference in tissue architecture, a rather similar mechanism appears to operate to build the apicolateral junctional belts. We show that the scaffold protein Nok/Mpp5 is necessary for the displacement of Pard3 and Cdh2 and for the formation of apical rings. In line with the *Drosophila* literature mentioned above, we propose that Nok operates here through its role in scaffolding Crumbs proteins to the apical surface. This proposal is consistent with the appearance of Crumbs protein at the nascent apical surface at the time when Pard3 and Cdh2 are displaced from this surface and the fact that Crumbs no longer localises to nascent apical surface in *nok* mutants (Figure 3C and Zou *et al*. 2013). A direct test of Crumbs function is difficult in zebrafish because three different Crumbs proteins (Crb1, Crb2a and Crb2b) are present at this stage in the developing neural rod. Previous research has shown that neuroepithelial organisation was only altered when all paralogs of Crumbs protein were knocked down by multiple morpholinos (Zou *et al*. 2013) and, consistent with our hypothesis, this appears to include a lack of lumen formation. Rather than transiently knocking down all paralogs of Crumbs proteins in this study, we used a Nok mutation (*nok*^m227^) that downregulates all paralogs of Crumbs protein from the nascent apical surface. In this case neither Pard3 nor Cdh2 were displaced from the nascent apical surface and this resulted in persistent adhesions between left and right sides of the neural rod and a loss of lumen formation. Additionally, we found that a stable genetic manipulation of Rab11a which also lacks Crumbs at the nascent apical surface (Roeth *et al*. 2009, Buckley et al 2013) also showed persistent contralateral adhesions between contralateral cells, a lack of lumen opening and, at later stages, comparable disorganisation of neuroepithelial tissue to *nok* mutants (Figure 4).

Despite its function in mediating epithelial cohesion via homophilic interactions of its extracellular domain (Thompson et al., 2013), the role of Crumbs protein in allowing cells to **resolve** their apical adhesions is starting to emerge as another important mechanism for controlling epithelial morphogenesis. For example, a recent study demonstrated that a secreted version of Crumbs2 acted non-cell autonomously to cause delamination of neuroepithelial cells during dorsal collapse of the spinal cord central canal in mice embryos (Tait et al., 2020). By studying dominant-negative Rab11a cells in an environment where they have access to wildtype neighbours, we uncovered that cells depleted of Crumbs from their apical surface can none-the-less form an epithelial surface when adjacent to wildtype cells. Thus, in the fish neuroepithelium, Rab11a and Crumbs appear to be required for *de novo* epithelial surface generation but dispensable for integration next to an already generated apical surface. This suggests that non cell-autonomous rescue from neighbouring wildtype cells is able to initiate epithelialisation in the absence of Rab11a/Crumbs. It would be interesting to determine whether secreted Crumbs protein plays a role in this rescue.

### Coordinating left-right release with epithelialisation

The zebrafish neural rod is initially a solid primordium, so the displacement of Pard3 and Cdh2 is not from a free apical surface but instead occurs within the densely packed cell-cell interfaces that lie at the centre of the rod. Cell adhesions between contralateral cells across the neural rod centre initially bind the left and right halves of the neural rod together, but these adhesions need to be released to generate the lumen and apical surface of the teleost neural tube. Our results show that Nok neatly coordinates the release of contralateral connections with the strengthening of ipsilateral connections, thus leading both to the generation of a free surface within the centre of an initially solid tissue and to junctional belt formation and epithelialisation. We found that Nok-deficient cells were unable to form a coherent sheet of apical rings (figure 1H). Additionally, in the absence of Nok or the presence of DNRab11a we found that cells were unable to release cell-cell adhesions. During epithelialisation of the neural rod, these defects first resulted in closed lumens (Figure 3B and 4Hi). This is in line with our previous findings in the neural tube (Buckley et al., 2013) and others’ recent findings that apical localisation of Rab11 is necessary for cell abscission and the normal opening of the Kupffer’s vesicle in zebrafish (Rathbun et al., 2020). It would be interesting to know whether similar mechanisms of junction remodelling also operate in the caudal segments of avian and mammalian spinal cord. The neural tube in these segments forms through the process of secondary neurulation that also involves the *de novo* formation of a central lumen within a condensing rod of cells (see for example Schoenwolf 1984 and Dady et al 2014), rather than the folding of epithelialized neural plate cells that generates the neural tube in the more rostral segments of these animals.

At later stages of development in the zebrafish neural tube, when neuroepithelial cells undergo interkinetic nuclear migration and D-divisions, the absence of Nok or the presence of DNRab11a resulted in progressive disintegration of midline organisation (Figure 4D and H). We propose that this disintegration is likely due to a combination of factors; during cell division, interkinetic nuclear migration and mitotic rounding pulls cells around within the tissue. In a normal epithelium, despite these forces, the apical surface would be stabilised by the lattice of ring-like junctional belts that hold the apical surface together. But when junctional belts between ipsilateral cells are not present, the tissue fails to hold cells at what should be the apical plane and cells are unable to fully re-extend to the basal side of the neural rod following division. Additionally, a lack of cell separation following division promotes the formation of local clumps of cells. Together, these defects cause divisions to occur in ectopic locations and the midline organisation disintegrates.

Our results suggest that an abnormal persistence of cell adhesions is a key part of the *nok* and DNRab11a phenotypes and this leads us to postulate that a lack of cell-cell separation may contribute to other reported epithelial disorganisation phenotypes in Crumbs or Rab11a-deficient epithelia (Das & Knust, 2018, Roeth et al., 2009).

### The role of the C-division

A recent paper also concluded that Nok is necessary to remodel junctional proteins during epithelialisation in the neural rod (Guo et al., 2018). They however propose that Nok’s role in apical remodelling is only necessary to rescue the tissue disruption caused by midline C-divisions, a conclusion based on their reported rescue of lumen formation and apical junctional belts when cell division is blocked in the *nok*^*m520*^ mutant. However, we show that blocking cell division does not rescue junctional belt formation in *nok*^*m227*^ mutants (Figure 7), and thus has a more fundamental role in apical junction remodelling than resolving the potential disruption to tissue organisation caused by C-divisions. We suggest the results of Guo et al (2018) may be a consequence of their use of the weaker *nok*^m520^ mutant allele having non-penetrant phenotypes, as they previously published (Zou et al., 2013). We further suggest the mini-lumen at the midbrain-hindbrain boundary that we observed in *nok*^m227^ mutants (Figure 6) is due to the expression of a second Nok-like gene expressed in this region (such as *mpp2b*, (Thisse & Thisse, 2004). *Nok/mpp5a* is a member of the Mpp MAGUK protein family and there are 6 *mpp* genes (MPP2-MPP7) with conserved protein domain structure in mammals. All of these genes contain a PDZ domain, which in the case of Nok (or Pals1), has been shown to be the specific domain that binds Crumbs (Roh et al., 2002). Little is known about the other *mpp* genes in zebrafish and this could form the basis of further study.

We have previously demonstrated that C-divisions at the midline of the neural rod are an important morphogenetic force during neural tube formation. Ectopic divisions are able to organise ectopic neural lumens and duplicate the neural tube (Tawk et al., 2007) while neural tubes generated in the absence of midline C-divisions are less efficient at resolving cell interdigitation across the midline and have a disorganised morphology (Buckley et al., 2013).

Here we show that connections between ipsilateral and contralateral cells are pulled into the cleavage furrow of midline C-divisions (Figure 2). This ingression of neighbours into the cleavage furrow is strikingly reminiscent of neighbour ingression during mitoses in *Drosophila* epithelia (Herszterg et al., 2013). Invasion of neighbours into the cleavage furrow also occurs during chick gastrulation where it can be developmentally regulated to promote or impede cell intercalation between the daughter cells and hence contribute to control of morphogenetic movements (Firmino et al., 2016). In the zebrafish neural rod, we show that multiple contralateral cell contacts are promoted through ingression of neighbouring cell processes into the cleavage furrow. We propose that the maintenance of connections with multiple contralateral cells through C-divisions ensures that cells do not solely connect to their sisters and we suggest this is one mechanism that leads to staggered connections across the midline. We suggest that this staggered alignment of endfeet may generate a morphogenetic advantage since it allows the attachment to a larger number of cells than a one-to-one ‘mirror adhesion’ as has previously been proposed (Guo et al., 2018), and therefore helps colocalise apical junctions from multiple neighbouring cells to the tissue midline. The localisation of junctional proteins at cell corners might also be important for allowing flexibility of cell movement during tissue remodelling (Finegan et al., 2019).

## Methods

### Embryo Care

All embryos were collected, staged and cultured according to standard protocols (Kimmel et al., 1995). All procedures were carried out with UK Home Office approval and were subject to local Ethical Committee review.

### Generation of Pard3-EGFP transgenic line

To generate the transgenic zebrafish line expressing Pard3-EGFP with endogenous spatial and temporal expression we used bacterial artificial chromosome (BAC) recombineering. We replaced the stop codon of the pard3-003/ASIP transcript (von Trotha et al., 2006), in the BAC clone DKEY-71E21 (Source Bioscience). This BAC clone contains ~71 kb upstream of the pard3-003/ASIP start codon and ~30 kb downstream from the targeted stop codon. We followed the protocol of (Bussmann & Schulte-Merker, 2011), with some modifications. The original protocol leaves a kanamycin selection cassette in the final BAC construct. However, we found this led to cell death and a lack of endogenous Pard3-EGFP expression. To remove the kanamycin selection cassette, we substituted two reagents with those from the Sarov *et al*., 2006 recombineering protocol and added an extra ‘flip out’ step to remove the kanamycin cassette. Specifically, we used the pRedFlp4 and R6k-EGFP-FRT-kanR-FRT cassettes from Sarov *et al* 2006. The EGFP-FRT-kan-FRT targeting cassette was PCR amplified with 50 bp homology arms at either end, to target it to replace the pard3-003/ASIP stop codon during BAC recombineering.

Forward primer: 5’- CACAGAAGCAGAACGGACGCAATGGACACCCCTCCACTTCAGACAGGTAC**AGCTCAGGAGGTAGCGG**-3’ and reverse primer: 5’ - AATTGAGTTTCATGATAGAACTTTGTATTTCTGCAATTCTGAAAAGCTGA**GGCAGATCGTCAGTCAG**-3’.

The protocol for pRedFlp4 transformation, targeted recombination steps and removal of the kanamycin resistance selection marker by pRedFlp4 induced flipase were carried out as previously described (Sarov et al., 2006), with slight modifications. Briefly, the recombineering pipeline was modified to have two rounds of tagging by Red/ET recombination, the first was insertion of iTol2_amp into the BAC backbone and the second insertion of EGFP-FRT-kanR-FRT. Successful recombination was assessed by colony PCR and then full sequencing of the sites of recombination. Preparation and injection of BAC DNA and screening for transgenic founders was all performed according to standard protocols (Bussmann & Schulte-Merker, 2011; Suster et al., 2011).

### Zebrafish Lines

The TgBAC(pard3:Pard3-EGFP)^kg301^ was generated as detailed above. The TgBAC(cdh2:Cdh2-tFT) (Revenu et al., 2014), was kindly provided by Darren Gilmour. The TgBAC(cdh2:Cdh2-tFT) line expresses both Cdh2-sfGFP and Cdh2-tagRFP, but we only image Cdh2-sfGFP expression in our experiments and refer to this as Cdh2-GFP for simplicity. These lines were bred with the Nok^m227^ mutant fish line (Wei & Malicki, 2002a) to generate compound Nok^m227^; Cdh2-tFT and Nok^m227^; Pard3-EGFP fish. To prevent ventricle opening in rhombomeres 3 and 5, we crossed the UAS-inducible dominant-negative Rab11a line, Tg(UAS:mCherry-Rab11a S25N)^mw35^ (Clark et al., 2011), with Tg(Krox20:RFP-KalTA4), which drives the optimised Gal4-activator, KalTA4, only in Krox20 positive rhombomeres 3 and 5 (Distel et al., 2009). This resulted in the expression of Rab11a-S25N specifically in rhombomeres 3 and 5 as previously described (Buckley et al., 2013).

### RNA preparation and injection

Fusion constructs containing cDNA in the pCS2+ vector were linearized and mRNA was synthesised using the SP6 mMessage mMachine kit (Ambion, AM1340). RNA for the following constructs was injected using standard protocols (Westerfield, 2000) at 40-100pg per embryo: Histone 2A tagged with GFP (H2A-GFP), Human Histone 2B tagged with RFP (H2B-RFP), Human CAAX membrane moiety tagged with EGFP, mCherry or mNeptune2.5 (EGFP-CAAX, mCherry-CAAX, mNeptune2.5-CAAX), Dominant-negative Human rab protein 11a tagged with EGFP (RAB11A-S25N-EGFP), Zebrafish partitioning-defective 3 tagged with GFP or RFP (Pard3-GFP/RFP). For ubiquitous distribution of mRNA, embryos were injected at the 1-2 cell stage. For mosaic labelling a single blastomere of a 16-64 cell stage embryo was injected.

### Morpholino injections

Co-injection of *nok* splice blocking morpholinos (donor: 5’-GTT TAT GAC ACC CAC CTA GTA AAG C -3’ and acceptor: 5’-CTC CAG CTC TGA AAG TAC AAA CAC A -3’), was made into one-cell stage embryos (Hsu et al., 2006). Full loss of Crb2a from the midline and associated phenotypes were seen with approximately 1.7 nL of 300 mM (0.5 pM) of each morpholino, which closely phenocopied the *nok* mutant phenotype (this manuscript supplementary figure 1 and Hsu *et al*. 2006). Mild to intermediate loss of Crb2a from the midline was seen with approximately 0.5-1 nL of 200-300 mM (0.1-0.3 pM) of each morpholino. The level of Crb2a loss correlated with the extent of associated phenotypes (see Supplementary figure 1). Standard morpholino sequence was 5’-CCT CTT ACC TCA GTT ACA ATT TATA 3’.

### Chemical inhibition of cell division

Cell division was blocked by incubating dechorionated embryos in 300 μM aphidicolin (Sigma) in 4% DMSO from bud stage (10 hpf) to 28 somite stage (23 hpf). Control embryos were incubated in 4% DMSO only. Embryos were maintained on a bed of agarose in 12-well plates in the dark throughout the drug incubation.

### Immunohistochemistry

Embryos were fixed with 4% paraformaldehyde for 2 hours at room temperature prior to processing for immunohistochemistry. The following primary antibodies were used: mouse anti-Crb2a (mouse monoclonal, ZIRC zs-4, 1:200), rabbit anti-Crb2a and anti-Crb1 (a kind gift from the Wei lab, 1:350), mouse anti-ZO-1 (mouse monoclonal, Invitrogen 33-9100, 1:500), rabbit anti-aPKC (rabbit polyclonal, Santa Cruz sc-216, 1:300), mouse anti-©tubulin (mouse monoclonal, MilliporeSigma T6557, 1:200) and mouse anti-ZO-1 (Life Technologies, 1:300). Rabbit anti-Pard3 (rabbit polyclonal, Millipore 07-330, 1:100) was used following fixation in Dent’s for 3 hours at room temperature and rehydration from methanol. AlexFluor secondary antibodies (Life Technologies) were used at 1:500 and Hoeshct (Life Technologies) nuclei stain was used at a final concentration of 1:10,000.

### Electron microscopy

Embryos were fixed in 2% paraformaldehyde and 1.5% glutaraldehyde in 0.1 M sodium cacodylate buffer over-night and then processed through 1% osmium and stained in 0.5% uranyl acetate. They were then dehydrated in ethanol and processed through propylene oxide into 100% resin, embedded and baked. Ultra-thin sections were taken in transverse orientation and imaged on a transmission electron microscope.

### Confocal imaging and processing

Embryos were mounted in low melting point agarose and imaged either using a Zeiss LSM 880 Fast Airyscan microscope and water dipping x20/1.0 N.A. objective, a Leica SP5 confocal microscope and water dipping x25/0.95 N.A objective or a PerkinElmer Ultraview spinning disk microscope. Imaging at the centre of a developing live tissue mass required the improved resolution in x, y and z obtained by using the Fast Airyscan mode of the Zeiss LSM 880 Fast Airyscan microscope. By using this microscope, high-resolution images of both *en face* Pard3-EGFP rings and the spatial location of Crb2a, aPKC and Cdh2-EGFP in the developing embryo were obtained. To further improve lateral imaging, fish were mounted on an agarose bed with small depressions to hold the yolk of the embryo in place. These agarose beds were made using a custom mould (a kind gift from Andrew Oates). Data was collected from the hindbrain and anterior spinal cord regions. Some adjacent cells have been removed from images of single cells to increase clarity of the cells of interest. Images were processed using Imaris, Volocity and Fiji/ImageJ.

### Data analysis

The custom R code used to generate plots for figures 1 and 3 is available at https://github.com/andyivanhoe/pard3-analysis.

To obtain apical ring density counts across the developing neuroepithelium (Figure 1E), Pard3-EGFP embryos were live-imaged en face between 15ss and 21ss using the Zeiss Airyscan. The dorso-ventral axis of the resulting z-stacks were divided into four quadrants (dorsal, mid-dorsal, mid-ventral, ventral), by applying a grid in Fiji. Apical rings were counted when all sides of the cell border had Pard3-EGFP signal along them.

To quantify the changing distribution of Pard3-EGFP across the neural rod midline over time we imaged 6 live TgBAC(pard3:Pard3-EGFP)^kg301^ embryos over a period of 3 hours using Zeiss Airyscan confocal acquisition and processing. We then made a maximum intensity projection of a 10 μm deep volume at a mid-dorsoventral level of the neural rod. A mediolateral line of 116 μm width was then drawn across the basal to basal width of the developing neuroepithelium and the mean fluorescence intensity at each pixel along that line measured using FIJI. This was repeated for 6 embryos at 1 hour intervals (Figure 3Aii, iii. 0 hours is 16 somite stage). All line profiles from different embryos and timepoints were aligned on the maximum intensity value for each line profile. The central 70 μm of the line profiles, centred on the maximum intensity values, were then used to calculate mean intensity and standard deviation at each point along the line for each timepoint before plotting. We aligned on maximum intensity value for each line profile because the exact maximum intensity of Pard3-EGFP relative to the tissue midline has slight biological variation between embryos.

To quantify the distribution of Pard3-EGFP at the midline at the fully neuroepithelial stage (after the single midline peak had resolved into what appeared to be two near parallel lines of expression on either side of the midline), we analysed a single z-level in the same 6 embryos where the two lines of expression were approximately 4 μm apart (Figure 3Aiv). This analysis was done at the last timepoint of the 3-hour timeline, but at a more ventral (and hence more morphogenetically advanced, see Figure 1 and associated text) position. In this case we measured intensity along a 23 μm wide mediolateral line. Because the midline expression is not perfectly straight and the peaks of expression only 4 μm apart, for this analysis we choose a single z-level and a thinner mediolateral line than for the timeline measurements to avoid averaging out the two peaks.

Plots in figure 4C, 4J, 7G and 7H were generated using Graphpad Prism. Specific statistical analyses are described in the figure legends.

## Acknowledgements

We thank Rachel Moore, Christopher Rookyard, Vineetha Vijayakumar and Xuan Liang for helpful discussions on this work. We thank Florent Campo-Paysaa for providing some of the movies analysed in figure 4H and 5B. We thank the Centre for ultrastructural imaging at KCL and the Cambridge Advanced Imaging for their help with processing electron microscopy samples. We thank the zebrafish facility in KCL for their help maintaining zebrafish stocks. We thank Darren Gilmour for the Cdh2:(Cdh2-tFT) line, Gwyn Gould for RAB11A-S25N-EGFP plasmid, Xiangyun Wei for the Crb2a antibody and Stefan Schulte-Merker and Mihail Sarov for BAC plasmid reagents. This work was supported by UK BBSRC grant BB/K000926/1 (JC), a Wellcome Trust Investigator Award (JC) and a Wellcome Trust/ Royal Society Sir Henry Dale Fellowship (CB).

## Author contributions

Conceptualization: AS, CB, JC

Investigation: AS, CB, CW, JC

Writing, reviewing & editing: AS, CB, JC

Supervision: CB, JC

Funding acquisition: CB, JC

