## Supplementary information for "Coordinated assembly and release of adhesions builds apical junctional belts during *de novo* polarisation of an epithelial tube"

**Supplementary movie 1 accompanies Figure 1G.**

This movie shows an example of the typical offset arrangement of apical rings either side of the tissue midline, prior to lumen opening. A Pard3-EGFP embryo was live-imaged *en-face* to the developing apical surface plane at 18-somite stage. The movie follows the z-stack from one side of the tissue midline, along the medio-lateral axis of the embryo, through the tissue midline to the other side of the embryo in 0.31 μm z-steps.

**Supplementary movies 2-4 accompany Figure 2A.**

Collectively, these movies demonstrate that the shape of the nascent apical endfeet (and therefore the location of the connection point between sister cells in relation to the rest of the cell shape) dynamically alters over the neural keel to rod transition. This is likely in response to the movements and mitoses of neighbouring (unlabelled) cells.

**Movie 2:** 8-minute-long movie illustrating the connection between sister cells from the 13-somite stage. The nascent apical endfoot of the left-hand cell transiently moves posteriorly during the movie.

**Movie 3:** 19-minute-long movie illustrating the connection between sister cells from the 14-somite stage. The connection point between sister cells is initially narrower than in the previous movie and then widens again to occupy the whole nascent apical endfeet of both cells.

**Movie 4:** 10-minute-long movie illustrating the connection between sister cells from the 15-somite stage. The nascent apical endfoot of the left-hand cell narrows and the relative position of its connection point with the right hand cell moves first to the anterior side and then to the posterior side of the right hand cell.

**Supplementary Figure 1: *nok* morphant and mutant comparison**

10µm Z-projections in dorsal orientation at a mid dorso-ventral level through the hindbrains of 30 h.p.f. embryos, stained via IHC for Crb2a and aPKC **i)** Wild-type embryos had open lumens lined apically with extensive Crb2a and aPKC protein (5/5 embryos). **ii)** In early *nok* morphant experiments, a lower concentration range of *nok* morpholinos was tested (approximately 0.1-0.3pM). This resulted in mild (17/21 embryos) or intermediate (11/21 embryos) loss of Crb2a from the tissue midline. The level of Crb2a loss correlated with the extent of associated phenotypes (lack of lumen opening, disorganised midline and ectopic apical proteins). **iii)** In later *nok* morphant experiments, a higher concentration of *nok* morpholinos was used (approximately 0.5pM). This resulted in an almost full or full loss of Crb2a from the posterior hindbrain tissue midline (3/3 embryos), a closed lumen, disorganised midline and ectopic apical proteins (6/8 embryos, see figure 4D). **iv)** *nok* mutant embryos phenocopied the later *nok* morpholino experiments, also showing a full loss of Crb2a from the posterior hindbrain tissue midline (4/4 embryos), a closed lumen, disorganised midline and ectopic apical proteins (9/9 embryos, see figure 4D).
