## Supplementary figures and images for "Coordinated assembly and release of adhesions builds apical junctional belts during *de novo* polarisation of an epithelial tube"

### Supplementary figure 1

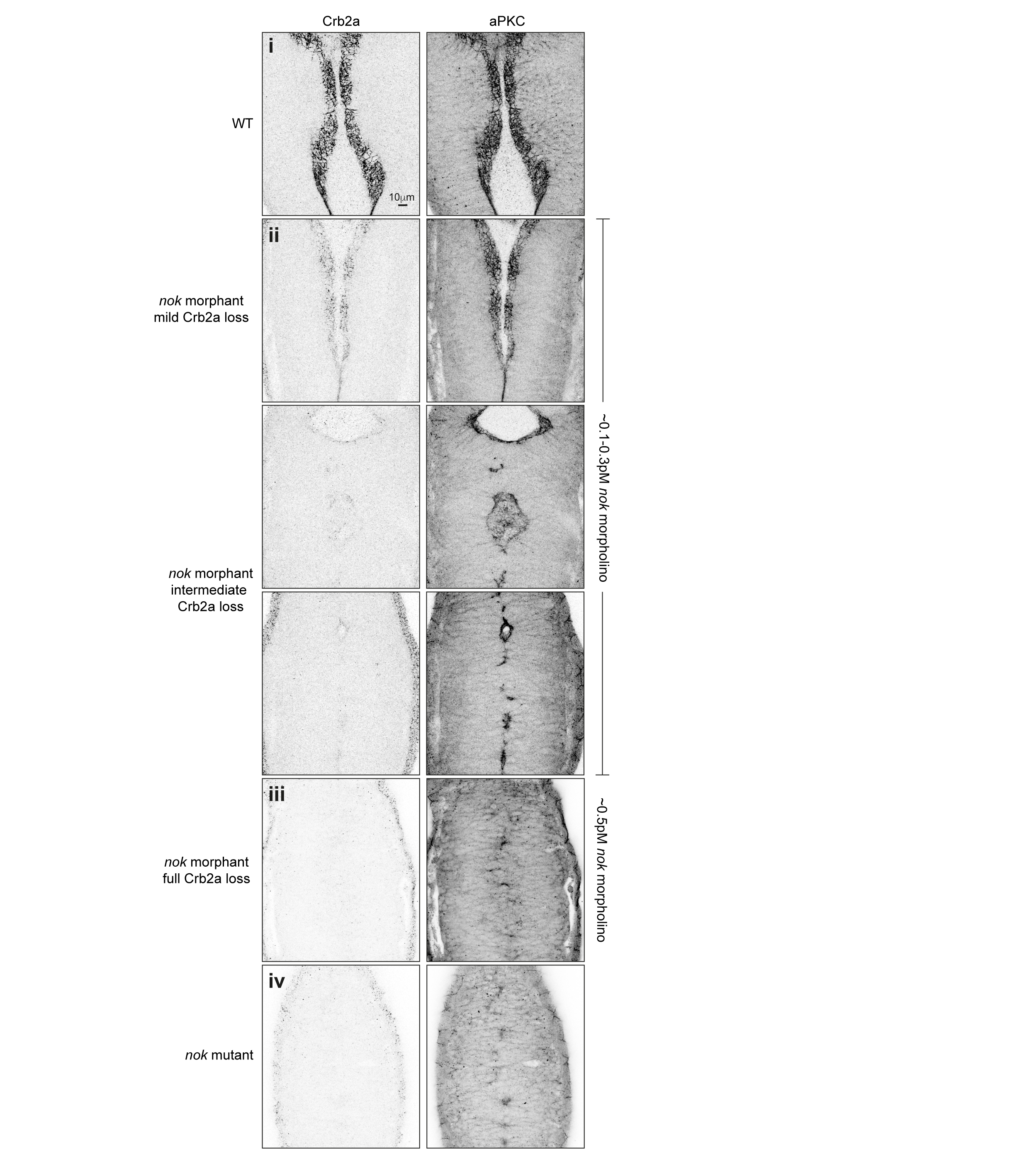
